# iDeLUCS: A deep learning interactive tool for alignment-free clustering of DNA sequences

**DOI:** 10.1101/2023.05.17.541163

**Authors:** Pablo Millán Arias, Kathleen A. Hill, Lila Kari

## Abstract

**Summary:** We present an *interactive* Deep Learning-based software tool for Unsupervised Clustering of DNA Sequences (*i*DeLUCS), that detects genomic signatures and uses them to cluster DNA sequences, without the need for sequence alignment or taxonomic identifiers. *i*DeLUCS is scalable and user-friendly: Its graphical user interface, with support for hardware acceleration, allows the practitioner to fine-tune the different hyper-parameters involved in the training process without requiring extensive knowledge of deep learning. The performance of *i*DeLUCS was evaluated on a diverse set of datasets: several real genomic datasets from organisms in kingdoms Animalia, Protista, Fungi, Bacteria, and Archaea, three datasets of viral genomes, a dataset of simulated metagenomic reads from microbial genomes, and multiple datasets of synthetic DNA sequences. The performance of *i*DeLUCS was compared to that of two classical clustering algorithms (*k*-means++ and GMM) and two clustering algorithms specialized in DNA sequences (MeShClust v3.0 and DeLUCS), using both intrinsic cluster evaluation metrics and external evaluation metrics. In terms of unsupervised clustering accuracy, *i*DeLUCS outperforms the two classical algorithms by an average of ∼ 20%, and the two specialized algorithms by an average of ∼ 12%, on the datasets of real DNA sequences analyzed. Overall, our results indicate that *i*DeLUCS is a robust clustering method suitable for the clustering of large and diverse datasets of unlabelled DNA sequences.

**Availability and implementation:** *i*DeLUCS is available at our github repository under the terms of the MIT licence.

**Supplementary information:** Supplementary data are available at *Bioinformatics* online.

## A Introduction

Clustering algorithms for DNA sequences play a fundamental role in bioinformatics, as they can be used to study the structural composition of DNA sequence datasets, to discover novel operational taxonomic units (OTU), and to complement phylogenetic analysis. The development of high throughput sequencing technologies has raised several challenges to many clustering methodologies, as most of them cannot keep up with the exponential increase in the number of sequences available for analysis. One of the reasons is that many clustering methods rely on the computationally expensive process of sequence alignment. To address these limitations, several alignment-assisted and alignment-free methodologies were proposed, see, e.g., [4, 2]. The majority of these methods also face scalability issues, as they are reliant on classic clustering algorithms that perform well in the low data regime but have poor performance when large amounts of data are available. While in the aforementioned approaches the exponential increase of data is a hindrance to good performance, other approaches, e.g., deep-learning-based methods, benefit from the availability of large amounts of data. In particular, multiple deep learning algorithms have been developed for classification and inference using both alignment-based and alignment-free methodologies [10, 9, 8]. It has also been shown recently by [7] that deep learning provides a significant improvement over classical unsupervised learning algorithms in discovering genomic-signature-based clusters, at different taxonomic levels. These promising initial results motivated the development of *i*DeLUCS, which takes advantage of the capabilities of deep learning, and is capable of clustering datasets comprising more than 400 Mbp. In addition, *i*DeLUCS exhibits several novel features which enhance the interpretability of its results: confidence scores of the final cluster assignments, a graphical user interface (GUI), dynamic visualization of the underlying training process, and incorporated evaluation metrics.

## B Software Description

*i*DeLUCS is a standalone software tool that exploits the capabilities of deep learning to cluster genomic sequences. It is agnostic to the data source, making it suitable for genomic sequences taken from any organism in any kingdom of life. *i*DeLUCS assigns a cluster identifier to every DNA sequence present in a dataset, while incorporating several built-in visualization tools that provide insights into the underlying training process and the composition of the datasets (Figure 1). *i*DeLUCS offers an evaluation mode to compare the ground-truth label assignments (or hypothesized label assignments) of the dataset sequences with their discovered cluster labels. This is accompanied by a visual qualitative assessment of the clustering, through the use of the uniform manifold approximation (UMAP, see [6]) of the learned lower dimensional embedding. Finally, *i*DeLUCS outputs confidence scores for all of its cluster-label predictions, for enhanced interpretability. The software was developed using Python 3.9 and can be deployed with or without a graphics processing unit (GPU) (see Appendix A for implementation details).

**Figure 1:**
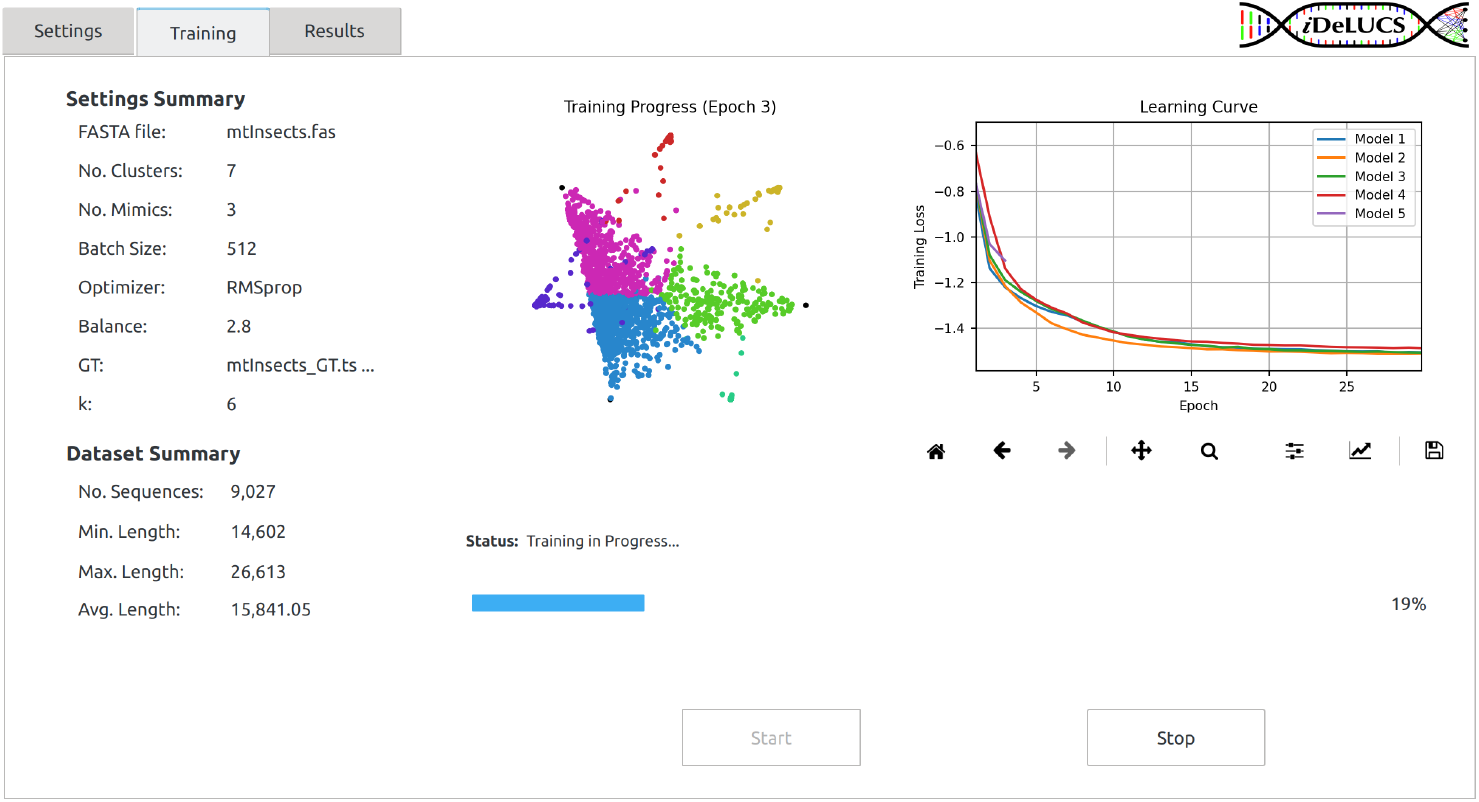
Training tab of *i*DeLUCS. The left panel displays a summary of the main training parameters, as well as some statistics about the dataset under study. The center panel contains a qualitative assessment of the learning progress. In this particular case, the figure illustrates the clustering of mitochondrial genomes of insects into 7 different clusters, each corresponding to one corner of the heptagon. Each point represents a genome, and its position indicates the probability that it is assigned to a different cluster/corner. The right panel contains a dynamic plot with the learning curves of the different models and serves as an indicator of whether or not the contrastive loss function is being minimized during training.

## C Materials and Methods

*i*DeLUCS builds upon the pipeline proposed in [7], consisting of: *(i)* calculating the *k*-mer frequencies for each DNA sequence, *(ii)* computing the data augmentations (mimic sequences), *(iii)* training multiple deep neural networks to learn the cluster assignments, and *(iv)* computing the majority voting cluster assignment for each sequence. In addition to multiple algorithmic optimizations to the pipeline, *i*DeLUCS significantly extends it in four main aspects. First, it uses the contrastive learning framework introduced by [1], and incorporates an additional contrastive term in the loss function, which enforces the consistency of the hidden representations learned by the artificial neural networks. These hidden representations are learned simultaneously with the cluster assignments via backpropagation. Second, it replaces the majority voting scheme by a more robust clustering ensemble based on information theory, which reduces the variance and boosts the accuracy. Third, it uses the information provided by the ensemble and the consistency of the hidden representations to provide an intrinsic quantitative assessment of the clustering assignment (silhouette coefficient, Davies-Bouldin Index), as well as to output the confidence score for the cluster assignment of each sequence in the dataset. Finally, the new contrastive learning framework can be combined with non-parametric clustering algorithms, such as HDBSCAN [5], to automatically determine the number of clusters. This *i*DeLUCS option is recommended only for fine-grained clusterings, due to the fact that HDBSCAN is a density-based method (see Appendix B for details).

To assess the performance and applicability of *i*DeLUCS, we first analyzed 14 real datasets with known ground-truth annotations (described in (a) and (b)): Nine datasets of mitochondrial DNA from various Kingdoms of life (Animalia, Protista, Fungi) totalling 18,810 sequences, two Bacteria datasets totalling 4,800 sequences, and three viral datasets totalling 4,144 sequences. In addition, we analyzed one dataset of simulated reads from microbial genomes (Bacteria and Archaea) comprising 432,333 reads (described in (c)), and 12 synthetic datasets totalling 246,625 artificial DNA sequences (described in (d)). Each dataset was selected for its unique characteristics, as described herein:

a. *Eight datasets from Kingdom Animalia, Kingdom Bacteria, and three datasets of viral sequences, obtained from [7]*: six mitochondrial DNA datasets of vertebrates at taxonomic levels from Subphylum to Family; two bacterial datasets to be clustered into families; and three viral datasets (Dengue, Influenza-A, Hepatitis B) to be clustered into virus subtypes. The maximum number of clusters per dataset is 12, and the maximum cluster size is 500 sequences, with the average sequence length of 16,700 bp for mtDNA, 433,882 bp for bacterial, and 5,058 bp for viral sequences.
b. *Three new mitochondrial DNA datasets* (Table 1) created to enhance the representation across Kingdoms of life: A dataset of 2,581 mitochondrial genomes from Kingdom Protista (average sequence length 17,141 bp) clustered into three phyla/subphyla; a dataset of 9,027 mitochondrial genomes from class Insecta (average sequence length 15,841 bp) clustered into seven orders; and a dataset of 1,759 mitochondrial genomes from Kingdom Fungi (average sequence length 62,644 bp), clustered into three phyla/subphyla.
c. *One dataset of simulated metagenomic reads from eight microbial genomes, obtained from [11]*. This dataset comprises more than 430,000 reads to be clustered into eight species (seven Bacteria and one Archaea). The reads were simulated using the PacBio sequencing simulation parameters, with maximum cluster size of 119,330 sequences, and average sequence length of 8,511 bp.
d. *12 synthetic datasets from [3]*. These are artificial datasets, each consisting of 100 random template sequences, representing the true clusters, and a random number of mutated copies that were generated from each template according to a pre-defined identity threshold. Each dataset contains at most 25,000 sequences, with a minimum dataset size of 18,210. The maximum number of clusters for each dataset is 12, the maximum cluster size is 400 sequences, and the average sequence length is 20,552 bp.

A detailed description of the datasets can be found in the Tables 1, 2, and 3 in Appendix C.

**Table 1:**
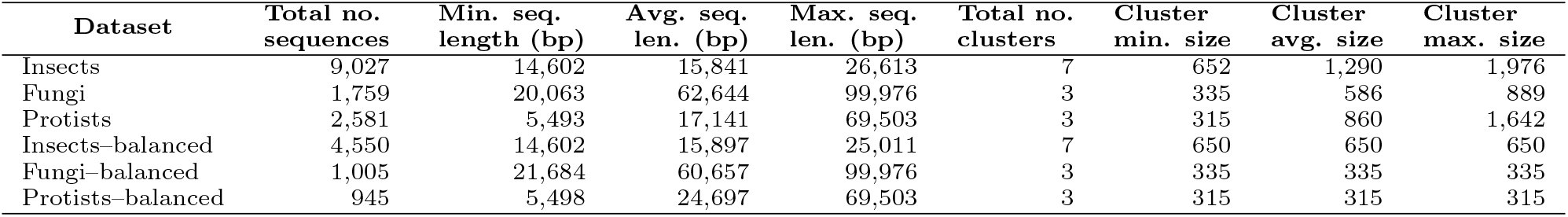
Description of the new mitochondrial DNA datasets (b). Note that there is a balanced version of each new dataset (Fungi, Protists, Insects). For the balanced version, the number of sequences per cluster was selected according to the number of sequences available in the smallest cluster.

**Table 2:**
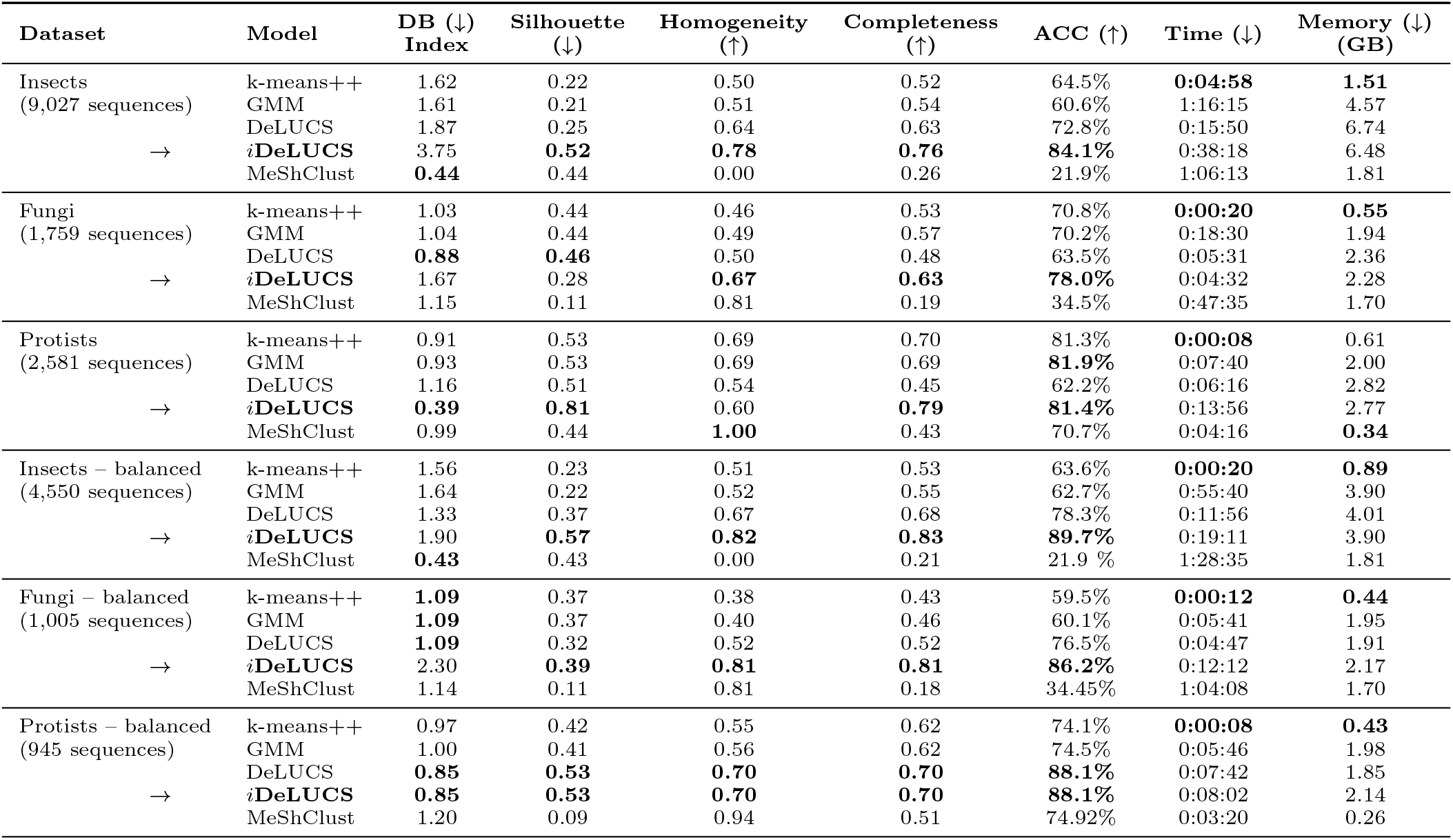
Comparison of the performance of *i*DeLUCS against *k*-means++, GMM, DeLUCS and MeShClust v3.0 clustering algorithms on the new mtDNA datasets (b), using intrinsic cluster evaluation metrics (Davies-Bouldin index, Silhouette coefficient) and external evaluation metrics (Homogeneity, Completeness, unsupervised accuracy ACC), as well as time and memory. Boldface indicates the best result, where (↑) indicates that higher is better, and (↓) indicates that lower is better. The top half of the table displays the results for the original datasets, and the bottom half those for the corresponding datasets with balanced clusters.

| Dataset | Model | DB (↓)<br>Index | Silhouette<br>(↓) | Homogeneity<br>(↑) | Completeness<br>(↑) | ACC (↑) | Time (↓) | Memory (↓)<br>(GB) |
| --- | --- | --- | --- | --- | --- | --- | --- | --- |
| Insects<br>(9,027 sequences) | k-means++ | 1.62 | 0.22 | 0.50 | 0.52 | 64.5% | **0:04:58** | **1.51** |
|  | GMM | 1.61 | 0.21 | 0.51 | 0.54 | 60.6% | 1:16:15 | 4.57 |
|  | DeLUCS | 1.87 | 0.25 | 0.64 | 0.63 | 72.8% | 0:15:50 | 6.74 |
|  | ***iDeLUCS*** | 3.75 | **0.52** | **0.78** | **0.76** | **84.1%** | 0:38:18 | 6.48 |
|  | MeShClust | **0.44** | 0.44 | 0.00 | 0.26 | 21.9% | 1:06:13 | 1.81 |
| Fungi<br>(1,759 sequences) | k-means++ | 1.03 | 0.44 | 0.46 | 0.53 | 70.8% | **0:00:20** | **0.55** |
|  | GMM | 1.04 | 0.44 | 0.49 | 0.57 | 70.2% | 0:18:30 | 1.94 |
|  | DeLUCS | **0.88** | **0.46** | 0.50 | 0.48 | 63.5% | 0:05:31 | 2.36 |
|  | ***iDeLUCS*** | 1.67 | 0.28 | **0.67** | **0.63** | **78.0%** | 0:04:32 | 2.28 |
|  | MeShClust | 1.15 | 0.11 | 0.81 | 0.19 | 34.5% | 0:47:35 | 1.70 |
| Protists<br>(2,581 sequences) | k-means++ | 0.91 | 0.53 | 0.69 | 0.70 | 81.3% | **0:00:08** | 0.61 |
|  | GMM | 0.93 | 0.53 | 0.69 | 0.69 | **81.9%** | 0:07:40 | 2.00 |
|  | DeLUCS | 1.16 | 0.51 | 0.54 | 0.45 | 62.2% | 0:06:16 | 2.82 |
|  | ***iDeLUCS*** | **0.39** | **0.81** | 0.60 | **0.79** | **81.4%** | 0:13:56 | 2.77 |
|  | MeShClust | 0.99 | 0.44 | **1.00** | 0.43 | 70.7% | 0:04:16 | **0.34** |
| Insects – balanced<br>(4,550 sequences) | k-means++ | 1.56 | 0.23 | 0.51 | 0.53 | 63.6% | **0:00:20** | **0.89** |
|  | GMM | 1.64 | 0.22 | 0.52 | 0.55 | 62.7% | 0:55:40 | 3.90 |
|  | DeLUCS | 1.33 | 0.37 | 0.67 | 0.68 | 78.3% | 0:11:56 | 4.01 |
|  | ***iDeLUCS*** | 1.90 | **0.57** | **0.82** | **0.83** | **89.7%** | 0:19:11 | 3.90 |
|  | MeShClust | **0.43** | 0.43 | 0.00 | 0.21 | 21.9 % | 1:28:35 | 1.81 |
| Fungi – balanced<br>(1,005 sequences) | k-means++ | **1.09** | 0.37 | 0.38 | 0.43 | 59.5% | **0:00:12** | **0.44** |
|  | GMM | **1.09** | 0.37 | 0.40 | 0.46 | 60.1% | 0:05:41 | 1.95 |
|  | DeLUCS | **1.09** | 0.32 | 0.52 | 0.52 | 76.5% | 0:04:47 | 1.91 |
|  | ***iDeLUCS*** | 2.30 | **0.39** | **0.81** | **0.81** | **86.2%** | 0:12:12 | 2.17 |
|  | MeShClust | 1.14 | 0.11 | 0.81 | 0.18 | 34.45% | 1:04:08 | 1.70 |
| Protists – balanced<br>(945 sequences) | k-means++ | 0.97 | 0.42 | 0.55 | 0.62 | 74.1% | **0:00:08** | **0.43** |
|  | GMM | 1.00 | 0.41 | 0.56 | 0.62 | 74.5% | 0:05:46 | 1.98 |
|  | DeLUCS | **0.85** | **0.53** | **0.70** | **0.70** | **88.1%** | 0:07:42 | 1.85 |
|  | ***iDeLUCS*** | **0.85** | **0.53** | **0.70** | **0.70** | **88.1%** | 0:08:02 | 2.14 |
|  | MeShClust | 1.20 | 0.09 | 0.94 | 0.51 | 74.92% | 0:03:20 | 0.26 |

To assess the performance of *i*DeLUCS on all the datasets analyzed in this study, we utilize both intrinsic (Davies-Bouldin Index, Silhouette Coefficient) and external (Homogeneity, Completeness, Unsupervised Clustering Accuracy (ACC)) clustering evaluation metrics. Of these, ACC arguably is the best indicator of performance, as it reflects the correspondence between cluster assignments and the ground-truth.

## D Applications and Results

The performance of *i*DeLUCS was compared against two classic clustering algorithms, *k*-means++ and Gaussian Mixture Models (GMM), as well as two recent clustering methods specific to DNA sequences, DeLUCS ([7]) and MeShClust v3.0 ([3]). The performance results for all four algorithms, in terms of intrinsic and external evaluation metrics as well as running time and memory usage, on the new mitochondrial datasets (b), are summarized in Table 2. The performance results for all algorithms on the other, previously published, datasets can be found in Tables S4, S5, S6 and S7 in Appendix C.

Overall, *i*DeLUCS has a robust performance across these very different types of datasets: small (113 sequences) or large (432,000 reads); real, simulated, or synthetic; at different taxonomic levels ranging from phyla to subtypes; with balanced clusters or with unbalanced clusters; with cluster number varying from 3 to 100 clusters; comprising long sequences (500,000 bp) or short sequences (650 bp); consisting of homologous sequences or of non-homologous sequences. On these datasets, the *i*DeLUCS clustering accuracy (ACC) ranges from 78% to 100%, with an average accuracy of 90%.

In particular, *i*DeLUCS outperforms the other four clustering algorithms on the real datasets in (a) and (b), most of which consist of non-homologous sequences. For example, for the mitochondrial genome datasets the average accuracy (ACC) of *i*DeLUCS is 92.4%, while classic clustering algorithms obtain an average accuracy of 70.2%, and the second best performant algorithm has an average accuracy of only 75.42%.

As seen in Table 2, for the new mitochondrial datasets (b), *i*DeLUCS obtains unsupervised classification accuracies ranging from 78% to 89.7%. Specifically, *i*DeLUCS outperforms all the other clustering algorithms for the balanced and unbalanced versions of the Insects and Fungi datasets, and has a comparable performance with the other classifiers for both the balanced and unbalanced versions of the Protist dataset. Note that the improved clustering ensemble and the new contrastive loss function of *i*DeLUCS significantly enhance its capability to cluster unbalanced datasets, compared to DeLUCS. These improvements become apparent in the clustering of the dataset of simulated long metagenomic reads (c), where the accuracy of *i*DeLUCS is ~ 16% higher than that of DeLUCS, and ~ 7% higher than that of *k*-means++.

For the synthetic datasets in (d), *i*DeLUCS obtains an average accuracy of 98.5% when the number of clusters is given as a parameter, and of 97.3% when the option of using HDBSCAN is selected to automatically determine the number of clusters. This is slightly lower than, but comparable to, the performance of MeShClust v3.0, which achieves an average accuracy of 99.3% for these synthetic datasets. Note that not all synthetic datasets analyzed in [3] were included in this comparison, since *i*DeLUCS was not optimized for some types of datasets. In particular, due to existing restrictions on the size of the output layer in deep learning models, the synthetic datasets where the expected number of clusters was large (5,000) were excluded. In addition, since the real genomic datasets analyzed in this study do not include short sequences (*<* 500 bp), the synthetic datasets with average sequence length *<* 500 bp were also excluded from this comparison. Future work is needed to systematically test and optimize *i*DeLUCS for datasets with short reads, or datasets where more than 200 clusters are expected.

All computational tests were performed on one of the nodes of the Beluga cluster of the Digital Research Alliance of Canada (2 x Intel Gold 6148 Skylake @ 2.4 GHz CPU, 32 GB RAM) with NVIDIA V100SXM2 (16GBh memory).

In summary, this study shows that *i*DeLUCS outperforms other algorithms in clustering sizeable datasets of unlabelled DNA sequences, especially when homology may or may not be present, and when the user has some prior knowledge of the expected number of clusters. Overall, our analysis shows that iDeLUCS is an accurate and scalable clustering method, performant on datasets of long, homology-free DNA sequences, not tractable via alignment-based methods due to either lack of alignment or excessive time complexity.

## Acknowledgements

The authors thank Shane Ding for his assistance with the testing and release of the software, Daniel Olteanu, Joseph Butler, and Connor Holmes for testing the software, and Zihao Wang for providing access to additional computational resources.

