## Supplementary Material for "iDeLUCS: A deep learning interactive tool for alignment-free clustering of DNA sequences"

### Supplementary Material
### "*i*DeLUCS: A deep learning interactive tool for alignment-free clustering of DNA sequences"

Pablo Millan Arias, Kathleen A. Hill, Lila Kari

The purpose of this document is to provide the user with all the information required to understand the functionality of the software tool *i*DeLUCS. The document is divided into three main parts: Appendix **A (Software Documentation)** contains information on how to use the tool and should be treated as a user manual, as it describes the training parameters, input formatting, data preparation, and the general functionality of the main tabs in the GUI. Appendix **B (Methodology)** describes the underlying theoretical principles behind *i*DeLUCS. Lastly, Appendix **C (Performance Evaluation)**, contains a description of the datasets used to benchmark the tool, the testing protocols and the results.

### A Software Documentation

#### A.1 Settings Tab

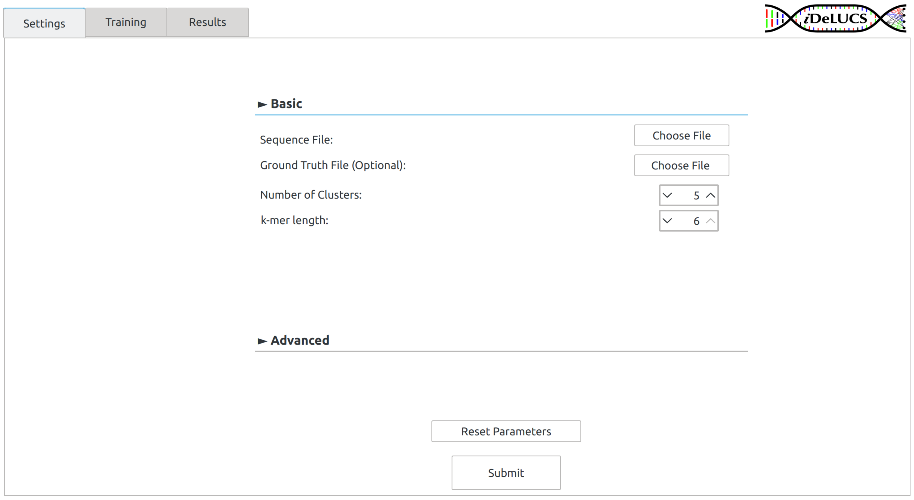

(a)

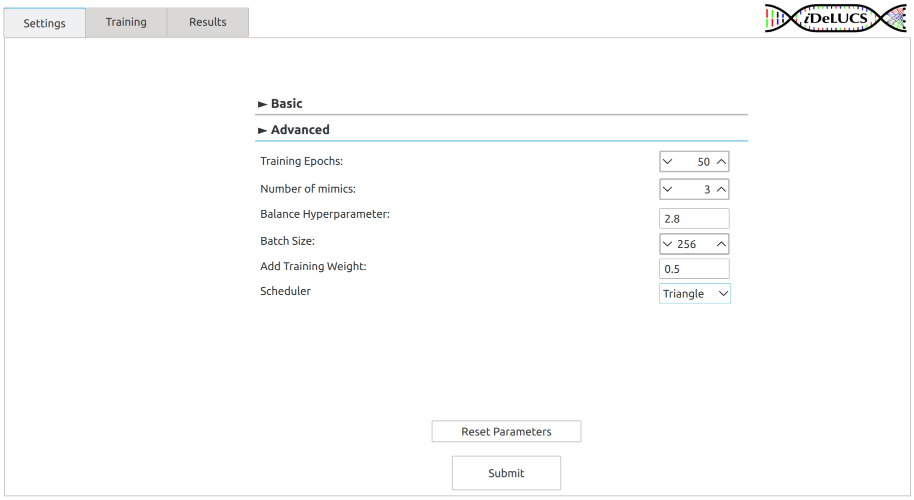

(b)

Figure 1: Two snapshots of the Settings Tab. (a) displays the "Basic" settings panel, where the users may select the parameters as they would for any classic clustering algorithm. (b) displays the "Advanced" settings panel, where the users can select all the training hyper-parameters specific to *i*DeLUCS.

This is the first interaction between the user and the tool, and an example of its configuration is illustrated in Figure 1. There are several hyper-parameters that are required for *i*DeLUCS to obtain the clustering assignment. The user may use the default values or select a specific value depending on the amount of information that is available about the dataset. All these parameters can be specified in the Settings tab of the program or through the command line. The following is a brief description of each parameter.

- `sequence_file`: *i*DeLUCS is multi-fasta format capable but not capable of multiple fasta files. All the sequences to be clustered must be joined into a single FASTA file and the path to this file must be provided as input in both the command line (CLI) and the graphical user interface (GUI) versions of *i*DeLUCS. The header of each sequence in the `sequence_file` must be a unique sequence identifier, and each sequence must follow the IUPAC nomenclature code for nucleic acids.

- `GT_file`: Tab-separated file with a hypothesized labelling for the training dataset, it must contain the following column names:

  - `sequence id` with the sequence identifiers, which must correspond to the identifiers provided in the headers of each sequence in the file.
  - `cluster_id` with the ground truth assignments for each sequence.

  Note that this is an optional parameter. The hypothesized labels will not be used during training, only for *post-hoc* analysis.

- `k`: k-mer length. (Default: 6; Options: {4, 5, 6}).

- `n_clusters` Expected or maximum number of clusters to find. This quantity should be greater or equal to `n_true_clusters` when `GT_file` is provided. (Default: 5; Range: [2,100]). The value of 0 is used for automatically finding fine-grained clusters through HDBSCAN.

- `n_epochs`: Number of training epochs. An epoch is defined as a training iteration over all the training pairs (*mimics*). (Default: 50; Range: [50,150]).

- `n_mimics`: Number of data augmentations per sequence that will be considered during training.(Default: 3; Range: [3,10]).

- `batch_sz`: Number of data pairs the network will receive simultaneously during training. A larger batch may improve convergence but it may harm the accuracy. (Default: 256; Range: [1,1024]).

- `lambda`: Hyper-parameter to control cluster balance. (Default: 2.8; Range: [1, 3]). Use a lower value for highly imbalanced datasets and a value closer to 3 when perfectly balanced clusters are expected.

- `weight`: Hyper-parameter to control the weight of the different loss functions. (Default: 0.25; Range: [0,1]). Use a higher value when low intra-cluster distance is expected and a lower value when high intra-cluster variability is expected.

- `scheduler`: Defines how the learning rate changes during training as detailed in [9]. (Default: `None` – Options:[`None`,`Triangle`, `Plateau`]).

##### A.2 Training Tab

In this tab users can monitor the status of the training process through qualitative information provided in the interactive plots (Figure 2). The left panel displays a summary of the main training parameters, as well as some statistics about the dataset under study. The center panel contains a dynamic figure representing how the assignments are evolving as training progresses. In this figure (central panel), each point represents a sequence, and its position indicates the probability that it is assigned to different clusters. Points that are located near the center have approximately equal probability to belong to all clusters, moreover, a point that is located near one of the vertices has a high probability of belonging to the cluster represented by that vertex and will be shown using the same color. The dynamic figure in the right panel of the tab, displays how the value of the contrastive loss changes as a function of the training epochs for each model used during training. These learning curves allow the user to visualize and assess the effects of the selected hyper-parameter on the training process.

##### A.3 Results Tab

This tab will be made available to the user either after the training is complete or after it was manually paused by the user (Figure 3). The left panel of the tab contains a table with the numeric cluster assignment for each sequence in the dataset, and a confidence score for each assignment. In the right panel, the tab will display a visual representation of the lower dimensional space that was learned by the network (model) with minimum loss during training. If the ground truth was provided, the tab will also display the confusion matrix calculated using the Hungarian matching algorithm, as well as the unsupervised clustering accuracy (ACC). If a ground truth is not provided, the Silhouette Coefficient and the Davies-Bouldin index, two other clustering metrics, are displayed for qualitative assessment of the clustering process. (See Appendix C.2 for a detailed description of the evaluation process)

##### A.4 *i*DeLUCS Output

After the user selects the 'Save Results' option on the Results Tab, or after the execution of the CLI version of the software is completed, a folder with the results will be provided as an output to the user. The name of the folder will

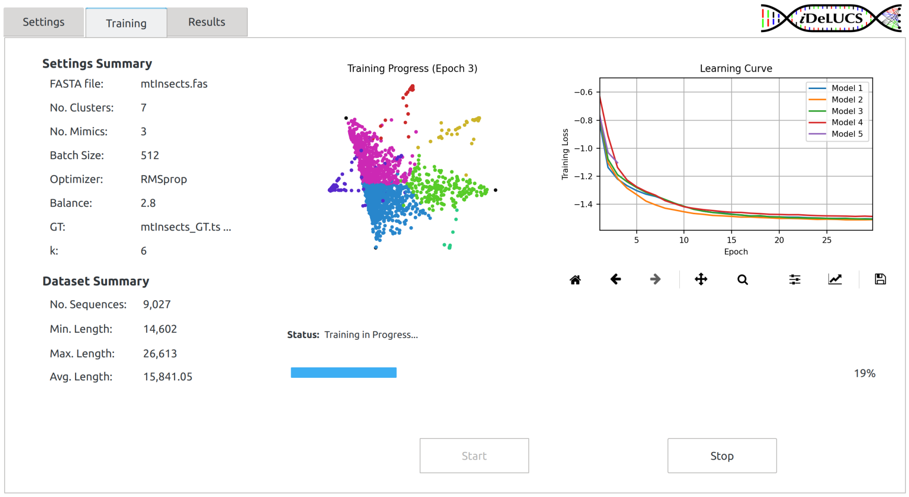

Figure 2: Snapshot of the training tab of $i$DeLUCS as it learns to cluster 9,027 mitochondrial genomes of insects into 7 different clusters. The left panel displays a summary of the main training parameters, as well as some statistics about the dataset under study. The center panel contains a qualitative assessment of the learning progress. The right panel contains a dynamic plot with the learning curves of the different models. Four models have been trained for thirty epochs each, and the training process of the fifth model is going through the third epoch.

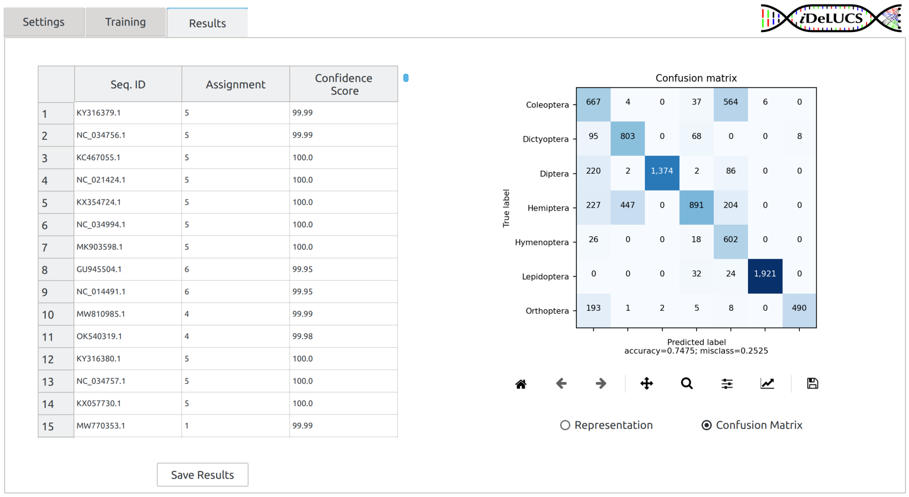

(a)

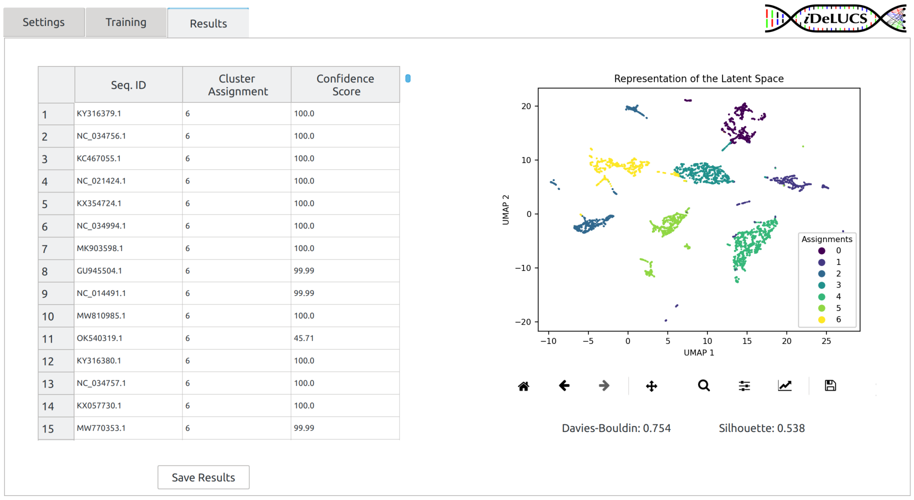

(b)

Figure 3: Snapshot of Results tab after the clustering process of 9,027 mitochondrial genomes of insects into 7 different clusters. Sub-figure (a) Displays the case where the ground truth labelling was provided, and the confusion matrix is calculated. Sub-figure (b) displays the case where no ground truth is provided, and only the lower dimensional representation and external clustering evaluation measures are provided instead.

be given according to the format: `Results/sequence_file_mmm_dd_hh:mm:ss`. The folder will contain the following files:

- `assignments.tsv`: File containing tabular data describing the cluster assignment of each sequence and the respective confidence score.
- `metrics.tsv`: File containing the value of the different clustering measures (ACC, Silhouette, Davies-Bouldin). When no ground truth is provided, only internal measures are calculated.
- `learned_representations.jpg`: Figure with the lower dimensional space that was learned by the network (model) with minimum loss during training. Similar to the right panel in Figure 3 (b).
- `training_plots.jpg`: Figure with the evolution of the contrastive loss during training for all models. Similar to the right panel in Figure 2.

##### A.5 Software Implementation

iDeLUCS is fully implemented in `Python 3.9`, and the source code is publicly available in the Github repository `https://github.com/Kari-Genomics-Lab/iDeLUCS`. For circumstances where the user only has access to a headless server, the CLI version of *i*DeLUCS provides similar functionality as the GUI version of the software. All the neural networks are implemented using `PyTorch 1.12` and the additional libraries are used in the software are: `numpy`, `pandas`, `sciki-learn`, `umap-learn`, `scipy` and `matplotlib`. For the CLI application, the training parameters hold the same identifiers as in the GUI but must be provided in a single line according to the following notation:

```
idelucs --sequence_file=<FASTA>
        --GT_file=<Optional Labelling>
        --n_clusters=3 --n_epochs=100
        --scheduler=None --lambda=2.8
        --batch_sz=512 --n_mimics=3
        --weight=0.25
```

#### B Methodology

##### B.1 The Contrastive Learning Framework

Various contrastive algorithms have been developed for unsupervised tasks such as clustering and representation learning. Although they are all different, most of these algorithms share the same principles, now known as the contrastive learning framework [2]. In this framework, which is at the core of *i*DeLUCS, there are three major components:

- *The construction of a dataset of paired data.* At this stage, given a data set $\mathcal{X} = \{x_1, \ldots, x_n\}$, the objective is to construct the set

$$X = \left\{ \left(\tilde{x}_1, \tilde{x}_1^1\right), \left(\tilde{x}_1, \tilde{x}_1^2\right), \left(\tilde{x}_1, \tilde{x}_1^3\right), \ldots, \left(\tilde{x}_i, \tilde{x}_i^m\right) \mid 1 \leq i \leq n \right\},$$

  where the data points in each pair $(\tilde{x}_i, \tilde{x}_i^j)$ are considered similar according to some criteria based on prior knowledge, e.g., invariance to distortions or spatial proximity. Note that samples $\tilde{x}_i$ and $\tilde{x}_i^j$ may not be present in the original dataset, and can be both an augmented version of the original sample $x_i$. This module is often referred to as the data augmentation module.

- *The definition of a neural encoder.* In order to learn from the paired dataset, the goal is to learn a mapping that encodes only what is common between $\tilde{x}_i$ and $\tilde{x}_i^j$, while dropping all the irrelevant information. If such a mapping $\Phi$ is found, the image $\mathcal{Y} = \Phi(\mathcal{X})$ becomes a lower dimensional representation of the original space $\mathcal{X}$ and can be used for downstream tasks. The best candidate for $\Phi$ is then a deep neural network $\Phi_\theta$ as its parameters $\theta$ can be optimized through backprogapation.

- *The contrastive loss function.* Depending on the unsupervised learning task of interest, any "pretext" task that attempts to minimize the distance between representations of pairs of samples can be used as inspiration for the loss function. A general form of a potentially successful contrastive loss is described in [1] as

$$\mathcal{L}_{\text{contrastive}} = w \cdot \mathcal{L}_{\text{alignment}} + (1 - w) \cdot \mathcal{L}_{\text{distribution}}$$

  where $\mathcal{L}_{\text{alignment}}$ encourages representations of paired samples to be consistent, while $\mathcal{L}_{\text{distribution}}$ encourages representations to match a target distribution. $w$ is the weighting parameter defining the importance of each term in the final loss.

#### B.2 *i*DeLUCS Pipeline

The methodology proposed in *i*DeLUCS builds upon the pipeline proposed in [6], in this subsection we illustrate how it fits into the contrastive learning framework. We describe its main components, and contrast them against the pipeline in [6].

##### B.2.1 Data Augmentation

The data augmentation module in [6] corresponds to the creation of $m$ artificial mimic sequences per original sequence using a probabilistic model based on transitions and transversions, while preserving the original sequences. The stochastic data augmentation model proposed in *i*DeLUCS transforms every DNA sequence in the dataset. This process produces two artificially created training samples, which are considered as positive training pairs even though they are not present in the original training dataset. We apply three different augmentations sequentially: Two random DNA substitution mutations, i.e., transitions and transversions with fixed independent substitution probabilities $p_{ts} = 10^{-3}$ and $p_{tv} = 5^{-3}$ respectively, and a random assignment of $r = 20$ nucleotides to symbol "N" (representing an unidentified nucleotide). This composition of augmentations incorporates robustness into the model and allows the networks to learn the structure of more complex datasets. Each augmented training sequence is then converted into a numerical vector containing the counts of all of its $k$-mers, where a $k$-mer is defined as a subsequence of length $k$ that does not contain the symbol "N". Finally, each $k$-mer count vector is converted into a k-mer frequency vector, via dividing its $k$-mer counts by the total length of the sequence minus the number of "N" symbols.

##### B.2.2 Neural Network - Base Encoder

We divide our architecture into a base encoder $f_\theta(\cdot)$, that extracts a meaningful lower dimensional representation, and a clustering layer $g_\theta(\cdot)$ such that $\Phi_\theta(x) = g_\theta(f_\theta(x))$. We employ the same architecture constructed in [6] as the backbone for $i$DeLUCS. For a mini-batch $X_{\mathcal{MB}} = (\mathbf{x}, \tilde{\mathbf{x}})$ of $2N$ augmented $k$-mer vectors, all the $k$-mer vectors are passed through the encoder, that consists of two fully connected layers, *Linear (512 neurons)* and *Linear (64 neurons)*, each one followed by a Rectified Linear Unit (*ReLU*) and a *Dropout* layer with dropout rate of 0.5. The hidden representation $\boldsymbol{z}_i = f(x_i)$ is then passed through the clustering layer *Linear (c_clusters)*, where $c$ is a numerical parameter representing the upper bound of the number of clusters, followed by a *Softmax* activation function.

##### B.2.3 Contrastive Loss Function

The pipeline in [6] uses the negative weighted mutual information as the loss function.

$$\mathcal{L}_I(\mathbf{x}, \tilde{\mathbf{x}}) = -\lambda \cdot H(\Phi(\mathbf{x})) + H(\Phi(\mathbf{x}) \mid \Phi(\tilde{\mathbf{x}})). \tag{1}$$

Note that Eq (1) fits the description of the general contrastive loss introduced in B.1. With the minimization of the loss function, the conditional entropy term $H(\Phi(\mathbf{x}) \mid \Phi(\tilde{\mathbf{x}}))$ should be as small as possible, resulting in sample $\mathbf{x}$ being perfectly predictable from $\tilde{\mathbf{x}}$. The entropy term can be interpreted as the Kullback-Leibler divergence between the output distribution and a uniform distribution over the cluster assignments $H(\Phi(\mathbf{x})) = KL(\Phi(\mathbf{x}) \mid\mid U(y))$, with $y \in \Phi(\mathcal{X})$. Furthermore, we observe that it is possible to learn the cluster assignments simultaneously with a hidden representation that allows the computation of distances between samples in different clusters. For this purpose, $i$DeLUCS combines the negative of the weighted mutual information with a loss function that enforces the consistency of the intermediate representations computed during training. We use the Normalized Temperature-scaled Cross

Entropy (NT-Xent) for this purpose:

$$\mathcal{L}_{\text{NT-Xent}} = -\frac{1}{n} \sum_{i,j<2N} \log \frac{\exp\left(\text{sim}\left(\boldsymbol{z}_i, \boldsymbol{z}_j\right)/\tau\right)}{\sum_{k=1}^{2n} \mathbf{1}_{[k \neq i]} \exp\left(\text{sim}\left(\boldsymbol{z}_i, \boldsymbol{z}_k\right)/\tau\right)} \tag{2}$$

Here, $\boldsymbol{z}_i = f(x_i)$, $\text{sim}\left(\boldsymbol{z}_i, \boldsymbol{z}_j\right) = \frac{\boldsymbol{z}_i \cdot \boldsymbol{z}_j}{\|\boldsymbol{z}_i\| \|\boldsymbol{z}_j\|}$ is the cosine distance, $\mathbf{1}_{[k \neq i]} \in \{0, 1\}$ is an indicator function evaluating to 1 iff $k \neq i$, and $\tau$ is a "temperature" parameter, used to restrict the range of the similarity scores, we use $\tau = 1$. The combined loss function in $i$DeLUCS can be written as:

$$\mathcal{L} = w \cdot \mathcal{L}_I + (1 - w) \cdot \mathcal{L}_{\text{NT-Xent}} \tag{3}$$

The hyper-parameter $w$ in Eq (3) controls the weight of the different loss functions. Figure 4 illustrates how $i$DeLUCS incorporates the additional contrastive term into the final loss to enforce the consistency of the hidden representations learned by the artificial neural networks. This provides robustness with respect to unbalanced datasets, as the learned representation of sequences in the same cluster are close to each other, but far from sequence in other clusters, see C.3 for more details.

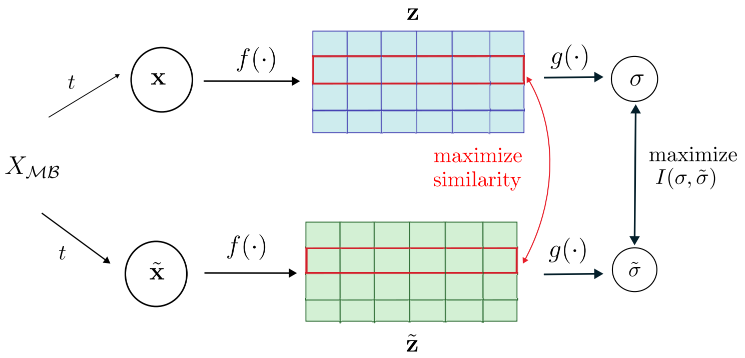

Figure 4: $i$DeLUCS maximizes the mutual information between the soft assignments $\boldsymbol{\sigma}$, $\tilde{\boldsymbol{\sigma}}$ of the augmentations $\mathbf{x}$, $\tilde{\mathbf{x}}$ from each training sequence after random mapping $t$, while maximizing the similarity of the hidden representations $\mathbf{z}$ and $\tilde{\mathbf{z}}$.

##### B.2.4 Information Theoretic Clustering Ensemble

For a given dataset $\mathcal{X} = \{x_1, \ldots, x_N\}$, a partition $\pi$ can be represented as a set of $K$ clusters $\pi = \{L_1, \ldots, L_K\}$, such that $\pi(x_i)$ denotes the cluster label assigned to $x_i$ by the partition. Suppose we are given a set $\Pi = \{\pi_1, ..., \pi_H\}$ of $H$ partitions of the data set $\mathcal{X}$The problem of clustering combination is to find the consensus partition $\pi_C$ that best summarizes the information present in $\Pi$. In general, the combination of multiple partitions in an unsupervised setting is a challenging problem, as each partition in the combination is represented as a set of labels assigned by an independent clustering algorithm with no trivial mapping between the assignments. Here we provide a detailed explanation of the concepts used in $i$DeLUCS which are a combination of the work presented in [7, 10, 11]. We use an information theoretic approach to the problem that does not compute an explicit mapping between the assignments. In this framework, the quality of the consensus partition $\pi_C = \{C_1, \ldots, C_K\}$ is determined by the amount of information it shares with all the partitions $\pi_i = \{L^i_j \mid 1 \leq j \leq K\} \in \Pi$. The best possible partition is then determined by

$$\pi_C^{\text{best}} = \arg\max_{\pi_C} \sum_{i}^{H} I(\pi_C, \pi_i) \quad \text{, where}$$

$$I(\pi_C, \pi_i) = \sum_{r=1}^{K}\sum_{r=1}^{K} p(C_r, L^i_j) \log\left(\frac{p(C_r, L^i_j)}{p(C_r)p(L^i_j)}\right)$$

is the classical Shannon mutual information between partitions. The previous optimization problem represents a difficult combinatorial problem [10]. However, the work in [11] shows that it is possible to consider a generalized definition of mutual information to simplify the problem. The generalized entropy of degree $s$ for a discrete probability distribution $P = (p_1, \ldots, p_n)$ is defined as

$$H^s(P) = (2^{1-s} - 1)^{-1}\left(\sum_{i=1}^{n} p_i^s - 1\right), \; s > 0, \; s = 1.$$

Hence, the generalized quadratic mutual information ($s = 2$) becomes:

$$\begin{aligned} I^2(\pi_C, \pi_i) &= H^2(\pi_i) - H^2(\pi_i \mid \pi_C) \\ &= -2\left(\sum_{j=1}^{K} p\left(L^i_j\right)^2 - 1\right) + 2\sum_{r=1}^{K} p\left(C_r\right)\left(\sum_{j=1}^{K} p\left(L^i_j \mid C_r\right)^2 - 1\right) \\ &= 2\sum_{r=1}^{K} p\left(C_r\right)\sum_{j=1}^{K} p\left(L^i_j \mid C_r\right)^2 - 2\sum_{j=1}^{K} p\left(L^i_j\right)^2 . \end{aligned} \tag{4}$$

With the following estimates, $p(C_r) = |C_r|/N$, $p(L^i_j) = |L^i_j|/N$, and $p(L^i_j \mid C_r) = |L^i_j \cap C_r|/|C_r|$, the quadratic mutual information $I^2(\pi_C, \pi_i)$ can be expressed in terms of the category utility score $\Delta$ as $I^2(\pi_C, \pi_i) = 2\Delta(\pi_C, \pi_i)$ [10]. This is relevant because in [7] Mirkin showed that a solution of the optimization problem of the utility function can be obtained through a transformation of the categorical labels into standardized binary features. $i$DeLUCS uses the same transformation and replaces each partition $\pi_i$ by $K$ binary features, and standardizes each binary feature to a zero mean. More specifically, for each data point $x$, and each partition $\pi_i \in \Pi$, the values of the new features are calculated as $y_{ij} = \mathbf{1}_{[L^i_j = \pi_i(x)]} - p(L^i_j)$, where $\mathbf{1}_{[x=y]} \in \{0, 1\}$ is an indicator function, evaluating to 1 if and only if $x = y$ . The final solution of the consensus partition problem can be obtained by a classic clustering algorithm operating over the new features $y_{ij}$. This clustering ensemble technique introduces robustness into the method and provides a better estimate of the confidence score of $i$DeLUCS for each sequence in the dataset.

##### B.3 iDeLUCS + HDBSCAN

It is possible that the number of expected clusters is unknown, in which case non-parametric clustering tools that automatically identify the number of clusters may be preferred. Fortunately, the contrastive learning framework can be seamlessly integrated with non-parametric clustering algorithms when some homology is expected. In this context, we have augmented $i$DeLUCS with an additional option to infer the number of clusters using the classical non-parametric clustering algorithm HDBSCAN [5]. To achieve this, we set the invariant information component of the loss in Equation 3, responsible for network assignment to zero, while the predominant component becomes the SimCLR loss. The learned 64-dimensional latent features are then used as input to HDBSCAN to compute the final clustering. We recommend using this feature only when the resulting clusters are expected to correspond to the lowest possible taxonomic level since HDBSCAN is a density-based method.

#### C Performance Evaluation

##### C.1 Sample Datasets

Besides the datasets provided in [6], users can test the performance of $i$DeLUCS over 3 additional mitochondrial datasets obtained from NCBI in June, 2022; one dataset of metagenomic reads simulated from eight microbial genomes using the Pacific Biosciences SMRT error model for long metagenomic reads, and 14 datasets of artificial DNA sequences created by first generating random template sequences and then mutating them by using various identity scores as provided by [4]. Dataset composition is summarized in Tables 1, 2, 3.

##### C.2 Evaluation Metrics

Clustering results can be evaluated *post hoc* using both internal and external validation methods, given the available information at test time. For the application domain of this paper (genomic datasets to be clustered according to their taxonomy), external evaluation methods seem more appropriate, as agreement with the ground truth (taxonomic groups) is ultimately a more important measure than other internal cluster properties such as separation or compactness. We select the unsupervised clustering accuracy (ACC) as the main external measure, as it utilizes the optimal mapping $f$ between discovered clusters $c_i \in C = \{c_i \mid i = 1, \ldots, Q\}$, and ground truth cluster labels $l_i \in L = \{l_i \mid i = 1, \ldots, K\}$, as provided by the Hungarian algorithm calculated using the contingency matrix $A = \{a_{ij}\}$ where $a_{ij}$ represents the number of

Table 1: Summary of the six mitochondrial DNA datasets of vertebrates, two bacterial datasets, and three viral datasets from [6] included in this study.

| Dataset | Total no. sequences. | Min. seq. len. (bp) | Avg. seq. len. (bp) | Max. seq. len. (bp) | Total no. clusters | Cluster min. size | Cluster avg. size | Cluster max. size |
|---|---|---|---|---|---|---|---|---|
| Vertebrata | 2,500 | 14,127 | 16,929 | 24,317 | 5 | 500 | 500 | 500 |
| Actinopterygii | 113 | 15,531 | 16,619 | 18,062 | 3 | 33 | 37 | 40 |
| Neopterygii | 1,475 | 15,468 | 16,687 | 19,763 | 6 | 226 | 245 | 250 |
| Ostariophysi | 363 | 15,664 | 16,635 | 17,998 | 3 | 120 | 121 | 123 |
| Cypriniformes | 545 | 16,061 | 16,608 | 17,280 | 7 | 70 | 77 | 85 |
| Cyprinidae | 447 | 16,070 | 16,632 | 17,426 | 12 | 26 | 37 | 47 |
| Bacteria | 3,200 | 150,499 | 433,614 | 500,000 | 8 | 400 | 400 | 400 |
| Proteobacteria | 1,600 | 150,499 | 434,150 | 500,000 | 4 | 400 | 400 | 400 |
| Dengue | 1,633 | 10,010 | 10,557 | 10,991 | 4 | 407 | 408 | 409 |
| HBV | 1,562 | 3,061 | 3,209 | 3,227 | 6 | 258 | 260 | 263 |
| Influenza-A | 949 | 1,345 | 1,409 | 1,468 | 5 | 187 | 190 | 193 |

Table 2: Summary of the three new mitochondrial DNA dataset and one dataset of simulated metagenomic reads from eight microbial genomes introduced by [12]. Note that there is a balanced version of each new dataset (Fungi, Protists, Insects), where the number of sequences per cluster in the balanced version was selected according to the number of sequences available in the smallest cluster.

| Dataset | Total no. sequences. | Min. seq. len. (bp) | Avg. seq. len. (bp) | Max. seq. len. (bp) | Total no. clusters | Cluster min. size | Cluster avg. size | Cluster max. size |
|---|---|---|---|---|---|---|---|---|
| Insects | 9,027 | 14,602 | 15,841 | 26,613 | 7 | 652 | 1,290 | 1,976 |
| Fungi | 1,759 | 20,063 | 62,644 | 99,976 | 3 | 335 | 586 | 889 |
| Protists | 2,581 | 5,493 | 17,141 | 69,503 | 3 | 315 | 860 | 1,642 |
| Insects (balanced) | 4,550 | 14,602 | 15,897 | 25,011 | 7 | 650 | 650 | 650 |
| Fungi (balanced) | 1,005 | 21,684 | 60,657 | 99,976 | 3 | 335 | 335 | 335 |
| Protists(balanced) | 945 | 5,498 | 24,697 | 69,503 | 3 | 315 | 315 | 315 |
| Simulated reads | 432,333 | 5,000 | 8,511 | 37,216 | 8 | 8,538 | 54,042 | 119,330 |

Table 3: Summary of the twelve synthetic datasets from [3] included in the study. The number in the name of each dataset represents an identity score threshold, indicating that each sequence in a cluster is within this threshold from the cluster center.

| Dataset | Total no. sequences. | Min. seq. len. (bp) | Avg. seq. len. (bp) | Max. seq. len. (bp) | Total no. clusters | Cluster min. size | Cluster avg. size | Cluster max. size |
|---|---|---|---|---|---|---|---|---|
| Medium-60 | 18,210 | 653 | 1,365 | 2,062 | 100 | 13 | 182 | 397 |
| Medium-70 | 18,731 | 678 | 1,359 | 2,027 | 100 | 7 | 187 | 399 |
| Medium-80 | 20,939 | 664 | 1,425 | 2,043 | 100 | 17 | 209 | 383 |
| Medium-90 | 21,266 | 730 | 1,340 | 2,016 | 100 | 9 | 213 | 398 |
| Medium-95 | 24,039 | 724 | 1,446 | 2,038 | 100 | 18 | 240 | 396 |
| Medium-97 | 20,772 | 736 | 1,358 | 2,022 | 100 | 6 | 208 | 399 |
| Long-60 | 20,885 | 1,393 | 2,758 | 4,039 | 100 | 10 | 209 | 400 |
| Long-70 | 18,558 | 1,441 | 2,754 | 4,062 | 100 | 6 | 186 | 399 |
| Long-80 | 20,525 | 1,396 | 2,639 | 3,974 | 100 | 9 | 205 | 396 |
| Long-90 | 22,518 | 1,489 | 2,586 | 3,964 | 100 | 6 | 225 | 397 |
| Long-95 | 20,222 | 1,461 | 2,890 | 4,049 | 100 | 14 | 202 | 401 |
| Long-97 | 19,960 | 1,486 | 2,715 | 3,988 | 100 | 9 | 200 | 396 |

sequences with label $l_i$ assigned to cluster $c_j$. The metric is formally defined as:

$$ACC = \frac{\sum_{i=1}^{n} O\left\{l_i = f\left(c_i\right)\right\}}{n}, \tag{5}$$

where $n$ is the total number of sequences, and for each DNA sequence $x_i, 1 \leq i \leq n$, we have that $f(c_i)$ is the taxonomic label assigned to $c_i$ by the optimal mapping $f$, and $O$ is a comparison operator returning 1 if the equality in the argument holds and 0 otherwise [6]. A value of 100% stands for a perfect match, and a value of 0% indicates that all samples were wrongly assigned. A larger value is correlated with a better matching with the ground truth. That being said, we also provide two additional external clustering metrics and two internal clustering metrics to complmenet the evaluation process and to provide a quantitative assessment of the performance of *i*DeLUCS for cases where the ground truth of the training data is not available.

- Homogeneity (higher is better): This external evaluation metric measures the extent to which each cluster contains only samples belonging to a single class [8]. It is defined as:

$$\text{homogeneity}(L, C) = \begin{cases} 1 & \text{if } H(L,C) = 0 \\ 1 - \frac{H(L|C)}{H(L)} & \text{otherwise} \end{cases}$$

  where $H(L|C)$ is the conditional entropy of the class distribution given the proposed clustering and $H(L)$ is the entropy of the true class labels.

The entropies are calculated as:

$$H(L|C) = -\sum_{c\in C}\sum_{l\in L}\frac{a_{lc}}{N}\log\frac{a_{lc}}{a_c}$$

$$H(L) = -\sum_{l\in L}\frac{n_l}{N}\log\frac{n_l}{N}$$

where $a_c$ is the number of sequences in cluster $c$ and $n_l$ is the number of sequences in true class $l$. Note that the homogeneity score ranges between 0 and 1, with 1 indicating perfect homogeneity. As noted by the authors in [8], in the perfectly homogeneous case, the value of $H(L|C) = 0$, when each cluster contains members of a single class.

- Completeness (higher is better): This external evaluation metric measures whether or not all data points that belong to a given class are assigned to the same cluster. In other words, a clustering result satisfies completeness if all data points from a single class are assigned to a single cluster. This metric, defined in [8] is symmetrical to homogeneity:

$$\text{Completeness}(L, C) = \begin{cases} 1 & \text{if } H(C, L) = 0 \\ 1 - \frac{H(C|L)}{H(C)} & \text{otherwise} \end{cases}$$

The entropies are calculated as:

$$H(L|C) = -\sum_{l\in L}\sum_{c\in C}\frac{a_{lc}}{N}\log\frac{a_{lc}}{n_l}$$

$$H(C) = -\sum_{c\in C}\frac{a_c}{N}\log\frac{a_c}{N}$$

where $a_c$ is the number of sequences in cluster $c$ and $n_l$ is the number of sequences in true class $l$. The completeness score ranges from 0 to 1, with higher values indicating better clustering performance.

- Silhouette Coefficient (higher is better): This measure compares the cluster assignment of a sequence with the assignment of the closest sequence assigned to a different cluster. Specifically,

$$Silhouette = \frac{1}{N}\sum_{i=1}^{N}\frac{b-a}{\max(a, b)}$$

where $N$ is the number of sequences in the dataset, $b$ is the distance between the representation of a sequence and the representation of the nearest sequence in a cluster it does not belong to, and $a$ is the mean intra-cluster distance. The best possible score is 1, and the score decreases as the overlap between the clusters increases. Scores with negative values indicate that most of the sequences have been placed in a wrong cluster, with the worst possible value being -1.

- Davies-Bouldin Score (lower is better): Defined as the average similarity measure of each cluster with its most similar cluster, it measures the average distance between clusters, relative to their sizes.

$$\textit{Davies-Bouldin} = \frac{1}{K}\sum_{i=1}^{K}\max_{i\neq j}\left(\frac{s_i + s_j}{d_{ij}}\right)$$

  where $s_i$ is the average distance between each point of cluster $i$ and the centroid $c_i$ and $d_{ij}$ is the distance between cluster centroids $c_i$ and $c_j$. Thus, assignment with high inter-cluster distance and low intra-cluster distance will result in a value closer to zero, which indicates a better score.

#### C.3 Performance Evaluation

To assess the performance *i*DeLUCS on the new datasets, we compare it against two classic clustering algorithms. The comparison is summarized in Table 2 and an example of the output confusion matrices is illustrated in Figure 5. Although *i*DeLUCS performs better overall on balanced datasets, both the improved clustering ensemble and the new contrastive loss function provide robustness with respect to unbalanced datasets, as Eq (3) is not dominated by the entropy term in favor of a uniform output distribution.
In addition, Table 3 summarizes the comparison between *i*DeLUCS and the pipeline proposed in [6] on eleven benchmarking datasets. We also explored specific improvements of *i*DeLUCS over the previous pipeline. In particular, we first explored the impact of the improved contrastive loss. For that purpose, we trained a single network using the loss function in Eq (3) over the new datasets. Figure 6 illustrates how enforcing the consistency of the hidden representations provides robustness with respect to unbalanced datasets, as the network learns an embedding where the representations of sequences in the same cluster are close to each other, but far from sequences in other clusters. We then compare the results against a single network trained using Eq (1) over the bechmarking datasets provided in [6]. Figure 7 shows how the newly introduced loss leads to better results both in terms of accuracy and convergence. Note that unlike the previous pipeline, *i*DeLUCS does not require the introduction of external noise on the parameters during training, as the model is less susceptible to getting trapped in local optima. Finally, we explored the impact of the clustering ensemble model introduced in *i*DeLUCS. For that purpose we trained a model with five voters, and compared the results of the final clustering ensemble with that of a majority voting scheme similar to the one used in [6]. Figure 8 shows how the newly introduced clustering ensemble produces better mean accuracy and lower variance for all the datasets.

**Note:** Reproducing Machine Learning experiments is a hard task. Completely reproducible results are not guaranteed across PyTorch releases, individual commits, or different platforms. Furthermore, results may not be reproducible between CPU and GPU executions, even when using identical seeds as it is said in PyTorch's official documentation. That being said, users may attempt to produce similar results as the ones obtained in this paper, by using the default parameters. Additional information about extra hyper-parameters and test scripts can be found on the paper repository. All of the tests were performed on one of the nodes of the Beluga cluster of the Digital Research Alliance of Canada (2 x Intel Gold 6148 Skylake @ 2.4 GHz CPU, 32 GB RAM) with NVIDIA V100SXM2 (16GB memory).

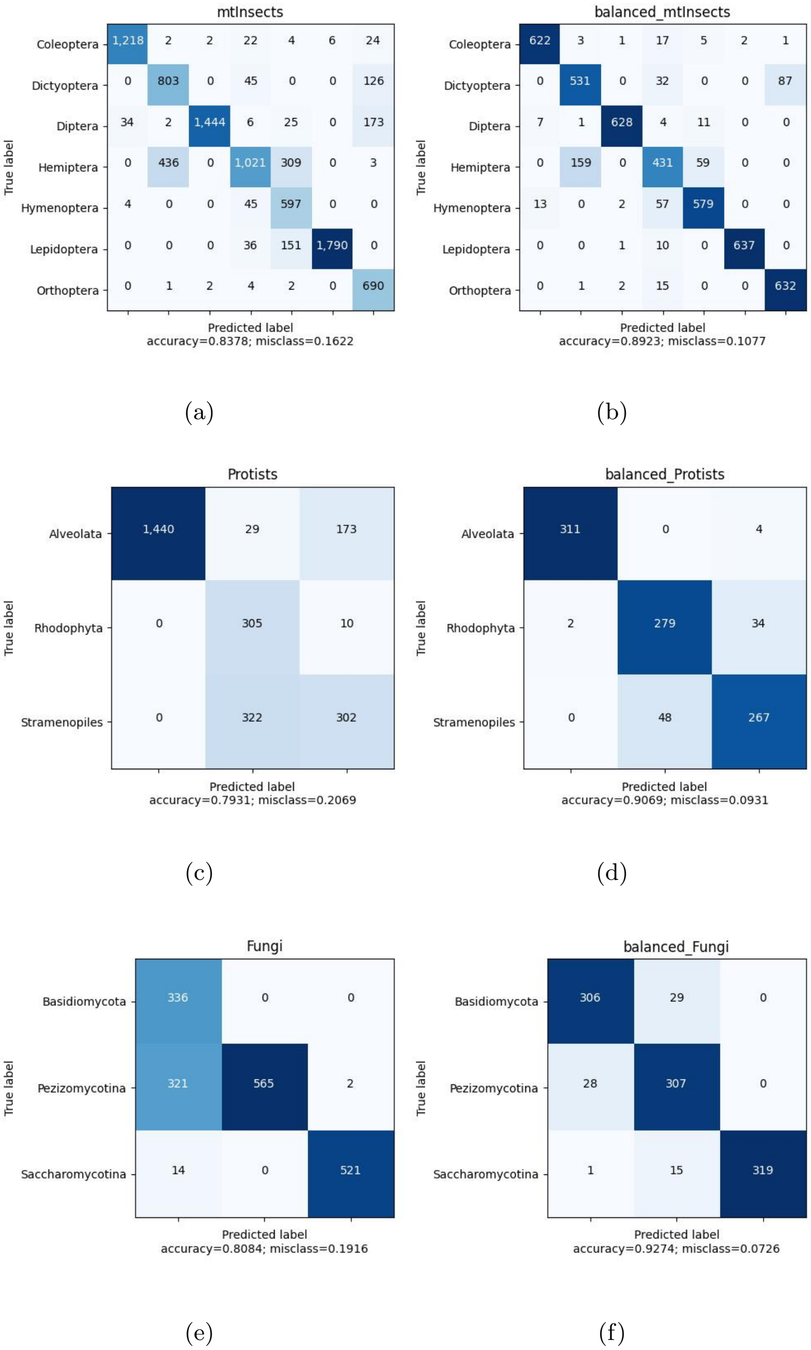

Figure 5: Confusion matrices with maximal accuracy obtained from the consensus clustering assignment and calculated via the Hungarian algorithm. The predicted labels are numeric cluster assignments, but are omitted for readability.

Table 4: Comparison of the performance of several clustering algorithms on the benchmark datasets introduced by [6], using both internal and external clustering evaluation measures. Boldface indicates the best result according to the convention (↑) higher is better and (↓) lower is better.

| Dataset | | Model | DB (↓) Index | Silhouette (↓) | Homogeneity (↑) | Completeness (↑) | ACC (↑) | Time (↓) | Memory (↓) (GB) |
|---|---|---|---|---|---|---|---|---|---|
| Vertebrata | | k-means++ | 2.65 | 0.11 | 0.65 | 0.67 | 81.3% | **00:00:17** | **0.65** |
| | | GMM | 2.96 | 0.09 | 0.63 | 0.71 | 67.7% | 00:29:44 | 2.90 |
| | | DeLUCS | 1.73 | 0.28 | 0.88 | 0.88 | 92.7% | 00:05:22 | 2.77 |
| | → | ***i*DeLUCS** | 0.96 | **0.56** | **0.98** | **0.98** | **99.4%** | 00:14:38 | 2.72 |
| | | MeshClust | **0.84** | -0.09 | 0.07 | 0.18 | 20.2% | 00:31:13 | **0.65** |
| Actinopterygii | | k-means++ | 1.87 | **0.26** | 0.86 | 0.86 | 94.69% | **00:00:8** | 0.32 |
| | | GMM | 2.55 | 0.16 | 0.41 | 0.51 | 65.48% | 00:02:25 | 1.82 |
| | | DeLUCS | **1.81** | 0.25 | **1.00** | **1.00** | **100.0%** | 00:00:59 | 1.71 |
| | → | ***i*DeLUCS** | 1.86 | **0.26** | **1.00** | **1.00** | **100.0%** | 00:01:58 | 1.77 |
| | | MeshClust | 0.95 | 0.15 | 0.52 | 0.56 | 65.5% | 00:00:31 | **0.18** |
| Neopterygii | | k-means++ | 2.01 | 0.20 | 0.59 | 0.66 | 64.3% | **00:00:07** | 0.52 |
| | | GMM | 3.10 | 0.14 | 0.52 | 0.56 | 62.5% | 01:33:36 | 3.01 |
| | | DeLUCS | 1.31 | **0.36** | 0.72 | 0.69 | 81.4% | 00:07:14 | 2.16 |
| | → | ***i*DeLUCS** | 2.54 | 0.31 | **0.73** | **0.73** | **86.0%** | 00:03:30 | 2.13 |
| | | MeshClust | **0.71** | 0.11 | 0.01 | 0.23 | 17.4% | 00:08:54 | **0.40** |
| Ostariophysi | | k-means++ | 4.01 | 0.06 | 0.63 | 0.66 | 84.3% | **00:00:39** | 0.35 |
| | | GMM | 3.44 | 0.07 | 0.64 | 0.67 | 82.6% | 00:18:07 | 1.82 |
| | | DeLUCS | 1.43 | **0.30** | 0.89 | 0.89 | 86.1% | 00:03:34 | 1.81 |
| | → | ***i*DeLUCS** | 2.10 | 0.22 | **0.96** | **0.96** | **97.2%** | 00:03:18 | 1.92 |
| | | MeshClust | **0.55** | 0.29 | 0.01 | 0.16 | 33.9% | 00:48 | **0.18** |
| Cypriniformes | | k-means++ | 2.65 | 0.12 | 0.66 | 0.68 | 75.8% | **00:00:03** | 0.39 |
| | | GMM | 2.62 | 0.12 | 0.67 | 0.68 | 75.8% | 00:13:22 | 3.47 |
| | | DeLUCS | 1.74 | 0.21 | 0.69 | 0.69 | 77.2% | 00:08:13 | 2.03 |
| | → | ***i*DeLUCS** | 1.01 | **0.45** | **0.73** | **0.74** | **80.7%** | 00:03:29 | 2.04 |
| | | MeshClust | **0.77** | 0.12 | 0.01 | 0.27 | 14.9% | 00:01:22 | **0.20** |
| Cyprinidae | | k-means++ | 1.96 | 0.28 | 0.90 | 0.90 | **91.1%** | **00:00:02** | 0.39 |
| | | GMM | 1.89 | 0.25 | 0.86 | 0.91 | 80.8% | 00:15:25 | 5.48 |
| | | DeLUCS | 1.96 | 0.33 | 0.90 | 0.89 | 90.6% | 00:07:29 | 1.89 |
| | → | ***i*DeLUCS** | **0.74** | **0.75** | **0.91** | **0.90** | **91.1%** | 00:08:59 | 1.97 |
| | | MeshClust | 0.92 | 0.05 | 0.42 | 0.46 | 11.9% | 00:00:47 | **0.19** |
| Bacteria | | k-means++ | 1.35 | 0.26 | 0.57 | 0.60 | 60.2% | **00:00:31** | **0.75** |
| | | GMM | 1.44 | 0.25 | 0.66 | 0.71 | 71.4% | 01:01:38 | 4.06 |
| | | DeLUCS | 1.42 | 0.36 | 0.71 | 0.71 | 79.1% | 00:12:05 | 3.17 |
| | → | ***i*DeLUCS** | 2.11 | **0.38** | **0.82** | **0.82** | **87.4%** | 0:21:54 | 3.14 |
| | | MeshClust | **0.88** | -0.23 | 0.38 | 0.29 | 20.0% | 10:42:35 | 8.19 |
| Proteobacteria | | k-means++ | 1.16 | 0.25 | 0.21 | 0.26 | 42.8% | **00:00:15** | **0.55** |
| | | GMM | 1.46 | 0.22 | 0.45 | 0.51 | 59.0% | 00:17:17 | 2.37 |
| | | DeLUCS | 1.22 | 0.28 | 0.75 | 0.76 | 89.9% | 00:09:38 | 2.25 |
| | → | ***i*DeLUCS** | 1.22 | **0.35** | **0.91** | **0.91** | **97.1%** | 00:14:08 | 2.20 |
| | | MeshClust | **0.96** | -0.22 | 0.24 | 0.18 | 29.2% | 02:27:33 | 4.72 |
| Dengue | | k-means++ | 1.02 | 0.44 | **1.00** | **1.00** | **100.0%** | **00:00:04** | **0.53** |
| | | GMM | 1.02 | 0.44 | **1.00** | **1.00** | **100.0%** | 00:04:00 | 2.33 |
| | | DeLUCS | 1.13 | **0.58** | **1.00** | **1.00** | **100.0%** | 00:01:49 | 2.25 |
| | → | ***i*DeLUCS** | 1.43 | 0.40 | **1.00** | **1.00** | **100.0%** | 00:02:42 | 2.22 |
| | | MeshClust | **0.68** | 0.28 | **1.00** | 0.50 | 51.2% | 00:02:14 | 0.19 |
| HBV | | k-means++ | 1.07 | 0.40 | **1.00** | **1.00** | **100.0%** | **00:00:04** | **0.52** |
| | | GMM | 1.35 | 0.33 | 0.87 | 0.93 | 75.6% | 00:09:50 | 3.13 |
| | | DeLUCS | **0.25** | **0.83** | **1.00** | **1.00** | **100.0%** | 00:01:48 | 2.19 |
| | → | ***i*DeLUCS** | 0.57 | 0.81 | **1.00** | **1.00** | **100.0%** | 00:02:41 | 2.17 |
| | | MeshClust | 0.62 | 0.28 | **1.00** | 0.81 | 93.3% | 00:00:40 | 0.15 |
| Influenza-A | | k-means++ | 1.26 | 0.40 | 0.98 | **0.98** | **99.4%** | **00:00:03** | 0.44 |
| | | GMM | 1.39 | 0.34 | 0.81 | 0.90 | 73.0% | 00:04:23 | 2.74 |
| | | DeLUCS | 1.39 | 0.31 | 0.98 | **0.98** | **99.4%** | 00:04:29 | 1.85 |
| | → | ***i*DeLUCS** | 0.86 | **0.58** | 0.98 | **0.98** | **99.5%** | 00:04:18 | 2.07 |
| | | MeshClust | **0.66** | 0.33 | **1.00** | 0.66 | 76.4% | 00:00:28 | **0.14** |

Table 5: Comparison of the performance of *i*DeLUCS against *k*-means++, GMM, DeLUCS and MeShClust v3.0 clustering algorithms on the new mtDNA datasets, using intrinsic cluster evaluation metrics (Davies-Bouldin index, Silhouette coefficient) and external evaluation metrics (Homogeneity, Completeness, unsupervised accuracy ACC), and Time, and Memory. Boldface indicates the best result, where (↑) indicates that higher is better, and (↓) indicates that lower is better. The top half of the table displays the results for the original datasets, and the bottom half those for the corresponding datasets with balanced clusters.

| **Dataset** | | **Model** | **DB (↓) Index** | **Silhouette (↓)** | **Homogeneity (↑)** | **Completeness (↑)** | **ACC (↑)** | **Time (↓)** | **Memory (↓) (GB)** |
|---|---|---|---|---|---|---|---|---|---|
| Insects | | k-means++ | 1.62 | 0.22 | 0.50 | 0.52 | 64.5% | **0:04:58** | **1.51** |
| | | GMM | 1.61 | 0.21 | 0.51 | 0.54 | 60.6% | 1:16:15 | 4.57 |
| | | DeLUCS | 1.87 | 0.25 | 0.64 | 0.63 | 72.8% | 0:15:50 | 6.74 |
| | → | ***i*DeLUCS** | 3.75 | **0.52** | **0.78** | **0.76** | **84.1%** | 0:38:18 | 6.48 |
| | | MeShClust | **0.44** | 0.44 | 0.00 | 0.26 | 21.9% | 1:06:13 | 1.81 |
| Fungi | | k-means++ | 1.03 | 0.44 | 0.46 | 0.53 | 70.8% | **0:00:20** | **0.55** |
| | | GMM | 1.04 | 0.44 | 0.49 | 0.57 | 70.2% | 0:18:30 | 1.94 |
| | | DeLUCS | **0.88** | **0.46** | 0.50 | 0.48 | 63.5% | 0:05:31 | 2.36 |
| | → | ***i*DeLUCS** | 1.67 | 0.28 | **0.67** | **0.63** | **78.0%** | 0:04:32 | 2.28 |
| | | MeShClust | 1.15 | 0.11 | 0.81 | 0.19 | 34.5% | 0:47:35 | 1.70 |
| Protists | | k-means++ | 0.91 | 0.53 | 0.69 | 0.70 | 81.3% | **0:00:08** | 0.61 |
| | | GMM | 0.93 | 0.53 | 0.69 | 0.69 | **81.9%** | 0:07:40 | 2.00 |
| | | DeLUCS | 1.16 | 0.51 | 0.54 | 0.45 | 62.2% | 0:06:16 | 2.82 |
| | → | ***i*DeLUCS** | **0.39** | **0.81** | 0.60 | **0.79** | **81.4%** | 0:13:56 | 2.77 |
| | | MeShClust | 0.99 | 0.44 | **1.00** | 0.43 | 70.7% | 0:04:16 | **0.34** |
| Insects – balanced | | k-means++ | 1.56 | 0.23 | 0.51 | 0.53 | 63.6% | **0:00:20** | **0.89** |
| | | GMM | 1.64 | 0.22 | 0.52 | 0.55 | 62.7% | 0:55:40 | 3.90 |
| | | DeLUCS | 1.33 | 0.37 | 0.67 | 0.68 | 78.3% | 0:11:56 | 4.01 |
| | → | ***i*DeLUCS** | 1.90 | **0.57** | **0.82** | **0.83** | **89.7%** | 0:19:11 | 3.90 |
| | | MeShClust | **0.43** | 0.43 | 0.00 | 0.21 | 21.9 % | 1:28:35 | 1.81 |
| Fungi – balanced | | k-means++ | **1.09** | 0.37 | 0.38 | 0.43 | 59.5% | **0:00:12** | **0.44** |
| | | GMM | **1.09** | 0.37 | 0.40 | 0.46 | 60.1% | 0:05:41 | 1.95 |
| | | DeLUCS | **1.09** | 0.32 | 0.52 | 0.52 | 76.5% | 0:04:47 | 1.91 |
| | → | ***i*DeLUCS** | 2.30 | **0.39** | **0.81** | **0.81** | **86.2%** | 0:12:12 | 2.17 |
| | | MeShClust | 1.14 | 0.11 | 0.81 | 0.18 | 34.45% | 1:04:08 | 1.70 |
| Protists – balanced | | k-means++ | 0.97 | 0.42 | 0.55 | 0.62 | 74.1% | **0:00:08** | **0.43** |
| | | GMM | 1.00 | 0.41 | 0.56 | 0.62 | 74.5% | 0:05:46 | 1.98 |
| | | DeLUCS | **0.85** | **0.53** | **0.70** | **0.70** | **88.1%** | 0:07:42 | 1.85 |
| | → | ***i*DeLUCS** | **0.85** | **0.53** | **0.70** | **0.70** | **88.1%** | 0:08:02 | 2.14 |
| | | MeShClust | 1.20 | 0.09 | 0.94 | 0.51 | 74.92% | 0:03:20 | 0.26 |

Table 6: Comparison of the performance of *i*DeLUCS against *k*-means++, DeLUCS and MeShClust v3.0 clustering algorithms on the dataset of simulated metagenomic reads from eight microbial genomes introduced by [12], using intrinsic cluster evaluation metrics (Davies-Bouldin index, Silhouette coefficient) and external evaluation metrics (Homogeneity, Completeness, unsupervised accuracy ACC), and Time, and Memory. Boldface indicates the best result, where (↑) indicates that higher is better, and (↓) indicates that lower is better. **Note:** GMM did not converge in this experiment.

| **Dataset** | | **Model** | **DB (↓) Index** | **Silhouette (↓)** | **Homogeneity (↑)** | **Completeness (↑)** | **ACC (↑)** | **Time (↓)** | **Memory (↓) (GB)** |
|---|---|---|---|---|---|---|---|---|---|
| Simulated reads | | k-means++ | 3.70 | 0.055 | 0.86 | 0.81 | 77.21% | **04:19:50** | **11.32** |
| | | DeLUCS | **0.56** | 0.68 | 0.65 | 0.70 | 67.34% | 06:27:45 | 17.34 |
| | → | ***i*DeLUCS** | 0.89 | **0.80** | **0.90** | **0.86** | **83.04%** | 05:25:30 | 17.50 |
| | | MeshClust | 4.12 | -0.24 | 0.0 | 0.0 | 0.0% | 33:45:07 | 16.00 |

Table 7: Comparison of the performance of $i$DeLUCS and $i$DeLUCS + HDBSCAN ($i$DeLUCS – auto) against $k$-means++ and MeShClust v3.0 clustering algorithms on the synthetic datasets introduced by [4], using intrinsic cluster evaluation metrics (Davies-Bouldin index, Silhouette coefficient) and external evaluation metrics (Homogeneity, Completeness, unsupervised accuracy ACC), and Time, and Memory. Boldface indicates the best result, where (↑) indicates that higher is better, and (↓) indicates that lower is better.

| **Dataset** | **Model** | **Clusters Found** | **DB (↓) Index** | **Silhouette (↓)** | **Homogeneity (↑)** | **Completeness (↑)** | **ACC (↑)** | **Time (↓)** | **Memory (↓) (GB)** |
|---|---|---|---|---|---|---|---|---|---|
| Medium_60 | k-means++ | 100 | 2.55 | 0.14 | 0.98 | 0.99 | 95.1% | **00:01:05** | 2.60 |
| | → $i$**DeLUCS** | 100 | 1.84 | 0.44 | **1.00** | 0.99 | 96.0% | 00:08:37 | 11.77 |
| | → $i$**DeLUCS** - auto | 68 | **0.86** | **0.64** | 0.93 | **1.00** | 90.3% | 00:40:38 | 11.78 |
| | MeShClust | 100 | 1.23 | 0.21 | **1.00** | **1.00** | **99.7%** | 00:24:41 | **5.35** |
| Medium_70 | k-means++ | 100 | 2.38 | 0.19 | 0.98 | 0.99 | 92.7% | **00:01:02** | 2.64 |
| | → $i$**DeLUCS** | 100 | 1.41 | 0.57 | **1.00** | 0.99 | 97.9% | 00:08:57 | 12.08 |
| | → $i$**DeLUCS** - auto | 79 | **0.74** | **0.72** | 0.97 | **1.00** | 94.1% | 00:42:36 | 12.08 |
| | MeshClust | 100 | 1.12 | 0.27 | **1.00** | **1.00** | **99.8%** | 00:22:56 | **5.66** |
| Medium_80 | k-means++ | 100 | 1.93 | 0.29 | 0.99 | 0.99 | 96.5% | **00:01:12** | 2.93 |
| | → $i$**DeLUCS** | 100 | **0.55** | **0.74** | **1.00** | **1.00** | **100.0%** | 00:10:11 | 13.35 |
| | → $i$**DeLUCS** - auto | 92 | 0.87 | 0.73 | 0.99 | **1.00** | 96.9% | 00:47:44 | 13.35 |
| | MeShClust | 100 | 0.89 | 0.35 | **1.00** | **1.00** | 99.8% | 00:25:55 | **7.02** |
| Medium_90 | k-means++ | 100 | 1.56 | 0.48 | 1.00 | 1.00 | 98.5% | **00:01:12** | 2.96 |
| | → $i$**DeLUCS** | 100 | 0.82 | **0.83** | **1.00** | **1.00** | 99.3% | 00:10:23 | 13.48 |
| | → $i$**DeLUCS** - auto | 100 | **0.59** | 0.87 | **1.00** | **1.00** | 99.9% | 00:48:53 | 13.48 |
| | MeShClust | 100 | 0.65 | 0.52 | **1.00** | **1.00** | **99.9%** | 00:20:52 | 7.24 |
| Medium_95 | k-means++ | 100 | 0.92 | 0.60 | 1.00 | 1.00 | 99.4% | **00:01:15** | 3.35 |
| | → $i$**DeLUCS** | 100 | 0.14 | **0.93** | **1.00** | **1.00** | **100.0%** | 00:11:38 | **15.07** |
| | → $i$**DeLUCS** - auto | 100 | **0.16** | **0.92** | **1.00** | **1.00** | **100.0%** | 01:41:49 | 15.14 |
| | MeShClust | 100 | 0.47 | 0.68 | **1.00** | **1.00** | **100.0%** | 00:22:01 | **9.20** |
| Medium_97 | k-means++ | 100 | 1.34 | 0.65 | 1.00 | 1.00 | 99.4% | **00:01:08** | 2.91 |
| | → $i$**DeLUCS** | 100 | 0.81 | 0.90 | **1.00** | 0.99 | 98.5% | 00:20:43 | 13.25 |
| | → $i$**DeLUCS** - auto | 100 | **0.11** | **0.96** | **1.00** | **1.00** | **100.0%** | 01:32:01 | 13.25 |
| | MeShClust | 100 | 0.37 | 0.74 | **1.00** | **1.00** | **100.0%** | 00:34:17 | **6.89** |
| Long_60 | k-means++ | 100 | 2.54 | 0.17 | 0.99 | 0.99 | 96.2% | **00:01:14** | 2.94 |
| | → $i$**DeLUCS** | 100 | 1.70 | 0.47 | **1.00** | 0.98 | 95.5% | 00:15:10 | 13.32 |
| | → $i$**DeLUCS** - auto | 76 | **0.81** | **0.67** | 0.96 | **1.00** | 90.8% | 01:24:05 | 13.32 |
| | MeShClust | 100 | 1.33 | 0.22 | **1.00** | **1.00** | **99.8%** | 00:47:43 | **7.04** |
| Long_70 | k-means++ | 100 | 2.24 | 0.23 | 0.98 | 0.99 | 96.4% | **00:01:02** | 2.64 |
| | → $i$**DeLUCS** | 100 | 1.61 | 0.56 | **1.00** | 0.99 | **97.0%** | 00:08:52 | 11.83 |
| | → $i$**DeLUCS** - auto | 82 | **0.47** | **0.74** | 0.98 | **1.00** | 96.6% | 00:38:08 | 11.98 |
| | MeShClust | 100 | 0.63 | 0.25 | **1.00** | 0.92 | 92.9% | 00:12:08 | **5.59** |
| Long_80 | k-means++ | 100 | 1.87 | 0.33 | 0.99 | 1.00 | 97.4% | **00:01:07** | 2.89 |
| | → $i$**DeLUCS** | 100 | 0.92 | **0.67** | **1.00** | **1.00** | 99.5% | 00:09:51 | 13.11 |
| | → $i$**DeLUCS** - auto | 100 | 0.92 | **0.67** | **1.00** | **1.00** | 99.5% | 00:38:51 | 13.11 |
| | MeShClust | 100 | **0.89** | 0.34 | **1.00** | **1.00** | **99.8%** | 00:26:39 | **6.79** |
| Long_90 | k-means++ | 100 | 1.07 | 0.51 | 1.00 | 1.00 | 99.1% | **00:01:14** | 3.16 |
| | → $i$**DeLUCS** | 100 | 0.53 | 0.82 | **1.00** | **1.00** | 99.6% | 00:10:47 | 14.26 |
| | → $i$**DeLUCS** - auto | 100 | **0.24** | **0.88** | **1.00** | **1.00** | **100.0%** | 00:41:31 | 14.26 |
| | MeShClust | 100 | 0.66 | 0.56 | **1.00** | **1.00** | **100.0%** | 00:21:45 | **8.14** |
| Long_95 | k-means++ | 100 | 0.71 | 0.62 | 1.00 | 1.00 | 99.4% | **00:01:03** | 2.86 |
| | → $i$**DeLUCS** | 100 | 0.72 | 0.88 | **1.00** | **1.00** | 98.8% | 00:15:52 | 12.94 |
| | → $i$**DeLUCS** - auto | 100 | **0.18** | **0.93** | **1.00** | **1.00** | **100.0%** | 00:35:46 | 12.94 |
| | MeShClust | 100 | 0.47 | 0.68 | **1.00** | **1.00** | **100.0%** | 00:11:06 | **6.60** |
| Long_97 | k-means++ | 100 | 0.98 | 0.69 | 1.00 | 1.00 | 99.8% | **00:01:04** | 2.86 |
| | → $i$**DeLUCS** | 100 | **0.10** | **0.95** | **1.00** | **1.00** | **100.0%** | 00:09:32 | 12.78 |
| | → $i$**DeLUCS** - auto | 100 | 0.12 | **0.95** | **1.00** | **1.00** | **100.0%** | 00:35:14 | 12.79 |
| | MeShClust | 100 | 0.38 | 0.73 | **1.00** | **1.00** | **100.0%** | 00:07:45 | **6.45** |

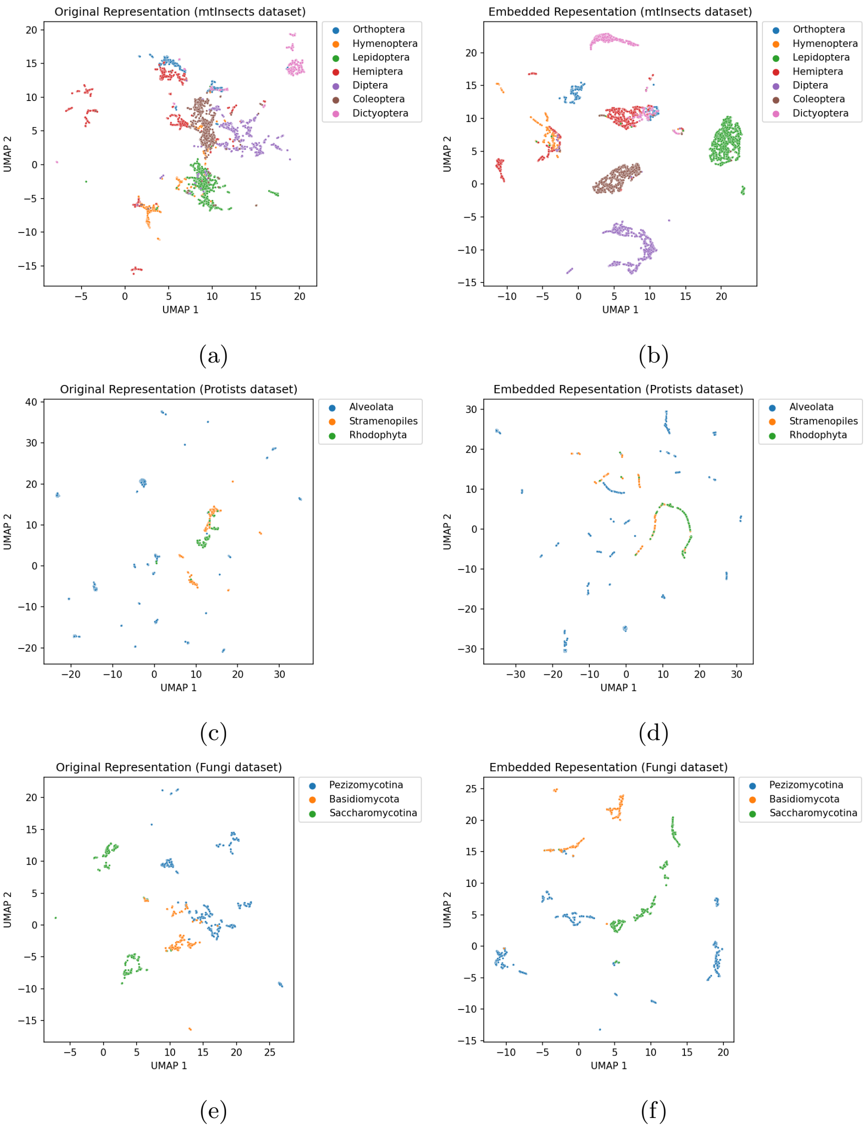

Figure 6: Comparison of the learned embedding with the original representation of the imbalanced newly introduced mtDNA datasets. $i$DeLUCS maps each sequence into a lower dimensional space that encodes the underlying taxonomy of the sequences for datasets with different compositions.

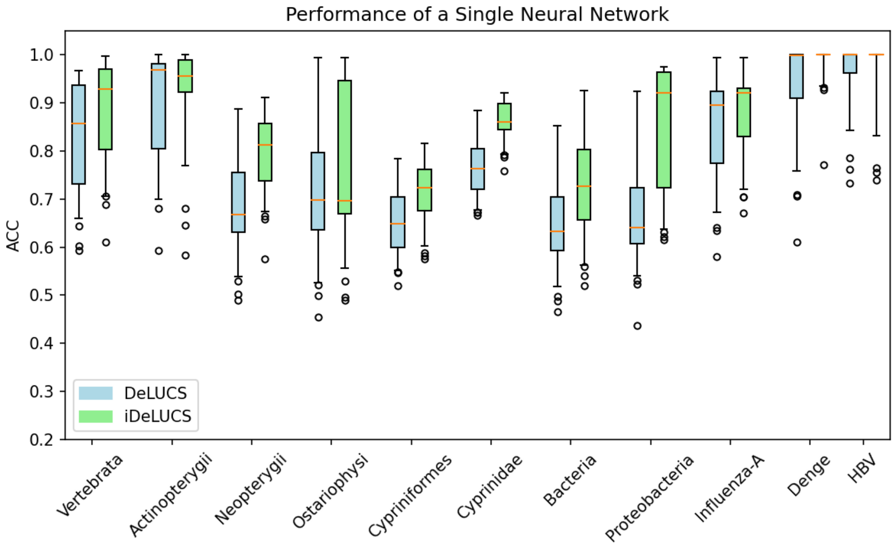

(a)

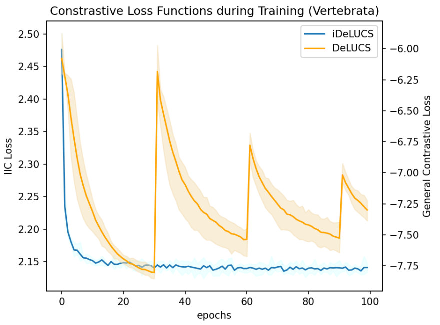

(b)

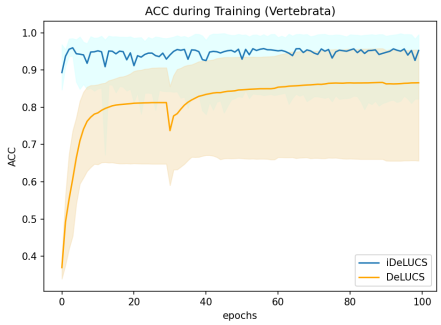

(c)

Figure 7: Comparison of the performance of $i$DeLUCS against the proposed pipeline in [6] on 11 benchmark datasets. (a) Box-plot representing the distribution of the unsupervised clustering accuracy produced by 100 neural networks trained independently over each dataset. Although, both DeLUCS and $i$DeLUCs produce an output with high variance, the mean ACC of the networks trained using the methodology of $i$DeLUCS is higher for ten out of eleven datasets. (b) Contrastive loss as a function of the training epoch for 100 runs of the training algorithm on the Vertebrata dataset. (c) Unsupervised clustering accuracy as a function of the training epoch for 100 runs of the training algorithm on the Vertebrata dataset.

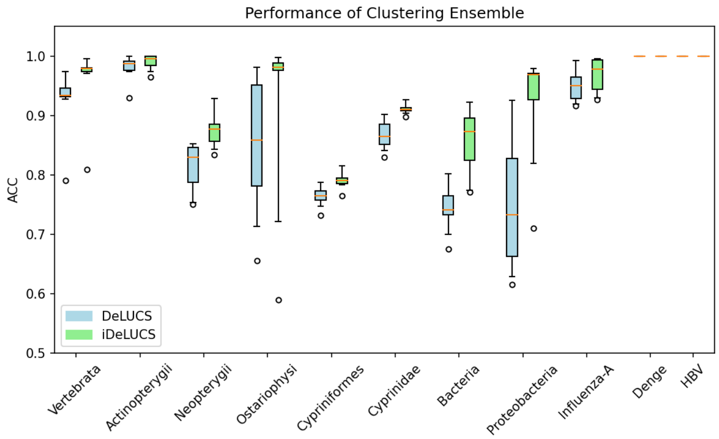

Figure 8: Box-plot representing the performance of the clustering ensemble of $i$DeLUCS against the majority voting in [6]. Fifty models with five voters were trained over the eleven benchmark datasets using both strategies. The clustering ensemble used by $i$DeLUCS outperforms the majority voting in [6] in all datasets.
